# Lifelong Cytomegalovirus and Early-Life Irradiation Synergistically Potentiate Age-Related Defects in Response to Vaccination and Infection

**DOI:** 10.1101/2021.12.01.470812

**Authors:** Jason L. Pugh, Christopher P. Coplen, Alona S. Sukhina, Jennifer L. Uhrlaub, Jose Padilla-Torres, Tomonori Hayashi, Janko Nikolich-Zugich

## Abstract

A popular “DNA-damage theory” of aging posits that unrepaired DNA damage leads to cellular (and organismal) senescence. Indeed, some hallmarks of immune aging are more prevalent in individuals exposed to Whole-Body Irradiation (WBI). To test this hypothesis in a model relevant to human immune aging, we examined separate and joint effects of lifelong latent Murine Cytomegalovirus (MCMV) and early-life WBI (i) over the course of the lifespan; (ii) in response to a West Nile virus (WNV) live attenuated vaccine; and (iii) following lethal WNV challenge subsequent to vaccination. We recently published that a single dose of non-lethal WBI in youth, on its own, was not sufficient to accelerate aging of the murine immune system despite causing widespread DNA damage and repopulation stress in hematopoietic cells. However, 4Gy sub-lethal WBI caused manifest reactivation of MCMV. Following vaccination and challenge with WNV in the old age, MCMV-infected animals experiencing 4Gy, but not lower, dose of sub-lethal WBI in youth had reduced survival. By contrast, old irradiated mice lacking MCMV and MCMV-infected, but not irradiated, mice were both protected to the same high level as the old non-irradiated, uninfected controls. Analysis of the quality and quantity of anti-WNV immunity showed that higher mortality in MCMV-positive WBI mice correlated with increased levels of MCMV-specific immune activation during WNV challenge. Moreover, we demonstrate that infection, including that by WNV, led to MCMV reactivation. Our data suggest that MCMV reactivation may be an important determinant of increased late-life mortality following early-life irradiation and late-life acute infection.

## INTRODUCTION

Susceptibility to infectious disease increases with age, making it one of the leading causes of death in people over 65 (1, 2). Aging is associated with a variety of immune defects affecting both innate and adaptive parts of immunity (rev. in (3)). Adaptive immunity age-associated defects include decreased naïve T cell numbers (4, 5), proliferation (6–9) and function (6–10) along with reduced pathogen-specific antibody production, somatic hypermutation and protective efficacy (11, 12). Theories of biological aging suggest that somatic cells, including immune cells, accumulate age-related defects, leading to impaired maintenance, function and, where applicable, selfrenewal; DNA damage-related senescence is one of the prominent factors implicated in these defects (13–17), but our understanding of long-term effects of DNA damage upon immune function in the old age in vivo is limited.

In addition, immune aging was hypothesized to be accelerated by the presence of life-long latent infections that impose a potential life-long burden on the immune system. One of the most prevalent latent viruses, the cytomegalovirus (CMV), has been associated in some [REF], but not other [REFS], studies to altered, suboptimal immune responses and increased all-cause mortality in both animal models and humans (18–24), although the mechanistic effects of CMV upon the aging immune system remain only partially understood. The most remarkable effect of CMV upon the immune system manifests in “memory inflation”– the presence of highly differentiated CMV-specific effector memory-phenotype T cells that increase in number throughout life (25, 26).

Whole-body gamma irradiation (WBI) results in systemic DNA damage (27) and hematopoietic lineage cell death in a dose-dependent manner (28–30). At WBI doses in the 0.5-4 Gray (Gy) range, hematopoietic cell populations are dramatically depleted. Barring severe host infection and death, in young organisms surviving immune cells divide rapidly and eventually repopulate to pre-irradiation levels (29). While the kinetics of immune repopulation has been described, long-term immune system performance following repopulation remains relatively uncharacterized. Hiroshima and Nagasaki bomb survivors display certain hallmarks of increased or premature immune aging (31–34). However, little is known about the influence of life-long CMV on the immunity of radiation survivors. In our previous study (Pugh, Aging Cell, 2016), we have shown that a single exposure of C57BL/6 (B6) mice to ionizing radiation of up to 4Gy in youth (equivalent to about 2Gy in humans) does not leave permanent scars on the immune system during aging, and that the animals responded well to vaccination and resisted subsequent challenge as well as their unirradiated counterparts.

However, experiments in specific pathogen-free animals are known to be misleading and there is an increasing realization that animal models should mimic common commensal and/or pathogen exposure in humans. To model the potential influence and interdependence of WBI, CMV, and aging on immunity, we here employed murine cytomegalovirus (MCMV) in a mouse model of WBI and natural aging. We tested vaccination and immunity in old age using the single-cycle live vaccine RepliVAX West Nile (R-WN) (35), followed by challenge with a potentially lethal dose of live West Nile Virus (WNV), as in our prior work (Pugh). Contrary to the results with uninfected irradiated mice, where vaccine and WNV-specific immunity, including T cell and antibody responses, were not substantially affected by WBI dose in youth, we found that in young adult animals carrying CMV, WBI induced immediate and clear reactivation of CMV. Mice carrying CMV and exposed to WBI in youth also exhibited signs of reduced immunity against CMV in the old age, as measured by reduced Th1 cytokine expression levels and percentages of cytokine-producing cells, increased expression of PD-1 on CMV-specific cells and reduced total anti-CMV antibody titers. In late life, CMV-positive animals irradiated in youth exhibited higher mortality following WNV challenge despite being vaccinated by R-WN, whereas animals exposed to WBI only or CMV only were fully protected, just as the control, unirradiated and uninfected animals. This reduced survival was associated with opportunistic MCMV reactivation during WNV challenge, likely resulting in a reduced ability of the irradiated immune system to deal with both the reactivated CMV and the WNV primary infection.

## RESULTS

### Hypotheses and Experimental Design

If DNA-damage related senescence is a causative factors in immune aging, we hypothesized that WBI would increase aging immune phenotypes in a dose-dependent manner. Further, if the effects of DNA damage are potentiated by cellular turnover in a causative manner, the combination of MCMV and WBI would be expected to result in an additive immune aging effect.

In order to test these hypotheses, we employed the experimental strategy outlined in **Fig. 1**. Age-matched, adult, male, C57BL/6 mice were divided into mock-infected and MCMV-infected groups at 2.3 months of age. Following a 60-day rest to allow for MCMV to establish stable latency, mice in each group were exposed to 0, 1, 2, or 4 Gray (Gy) WBI in a single dose. As would be expected from the WBI LD50/30 of this strain (37), no mice died from the exposure. 72 hours post WBI, as well as at 13 and 19 mo of age, 6 mice/group were analyzed cross-sectionally for lymphoid cell depletion/death and repopulation in a WBI dose-dependent manner. In addition, immune populations in peripheral blood were tracked at 3-month intervals from WBI until 19 months of age. At 19 months of age, mice were injected with RWN vaccine, which we have previously shown to be protective from WNV in old mice (36). Following development of primary and memory immune responses (60 days post infection), we challenged mice with a potentially lethal dose of live WNV and determined survival. Following both vaccination (day 7) and WNV challenge (day 67), we also measured WNV-specific adaptive immunity as described below. This design was replicated on two independent cohorts of mice separated by approximately one year, with comparable results.

**Figure 1:**
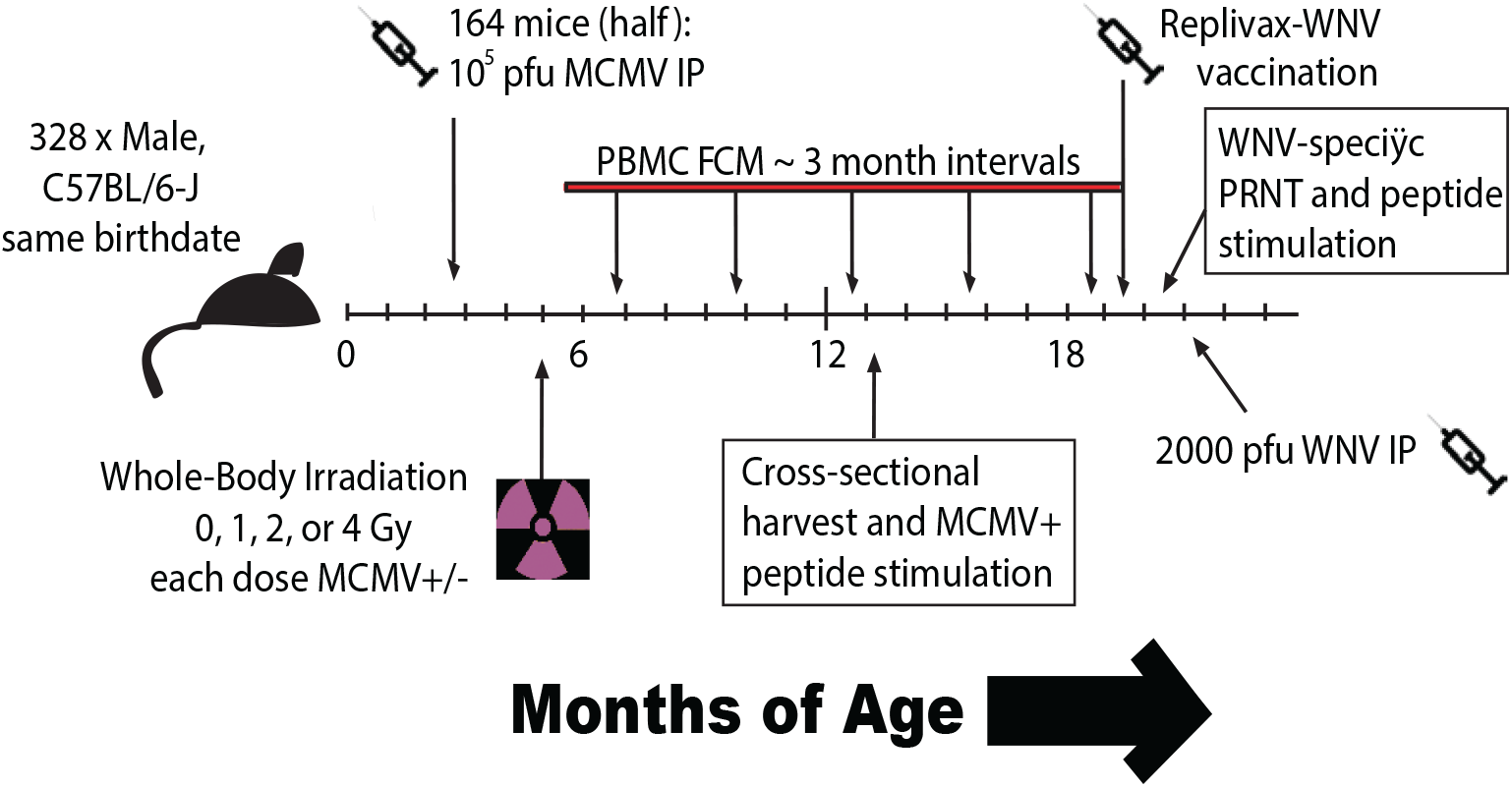
Experimental design of longitudinal cohorts. Age-matched, adult, male, C57BL/6 mice were divided into those receiving MCMV(+), and those remaining uninfected with MCMV(-), for life. Following a 60-day rest, mice in each group were then further divided into those receiving 0, 1, 2, or 4 Gray (Gy) of WBI in a single dose. A cross-sectional harvest of representative mice from each group was collected at 13 months of age, long after complete repopulation of immune cells. In addition, immune populations in peripheral blood were tracked at 3-month intervals from WBI until 19 months of age. At 19 months of age, mice were injected with 10^5^ pfu RWN vaccine IP. Immune function and antibody generation were assayed 45-days post-vaccination. 60 days post-vaccination, mice were challenged with 2000 pfu WNV IP.

### High-dose WBI in youth results in reduced survival from WNV challenge in old age only in mice with life-long MCMV

Following vaccination and WNV challenge, the vast majority of mice in the MCMV(+) 0Gy, MCMV(-) 4Gy and MCMV (-) 0Gy (control) groups all survived WNV challenge. However, survival in MCMV(+) 4Gy mice was significantly worse compared to either of the above three groups (**Fig. 2**). Lower irradiation doses (1 & 2 Gy) had no significant impact on survival regardless of the presence of MCMV (**Fig. S1**). Therefore, neither WBI alone nor CMV alone, administered in youth, could adversely affect the ability of an old organism to survive WNV challenge following vaccination.

**Figure 2:**
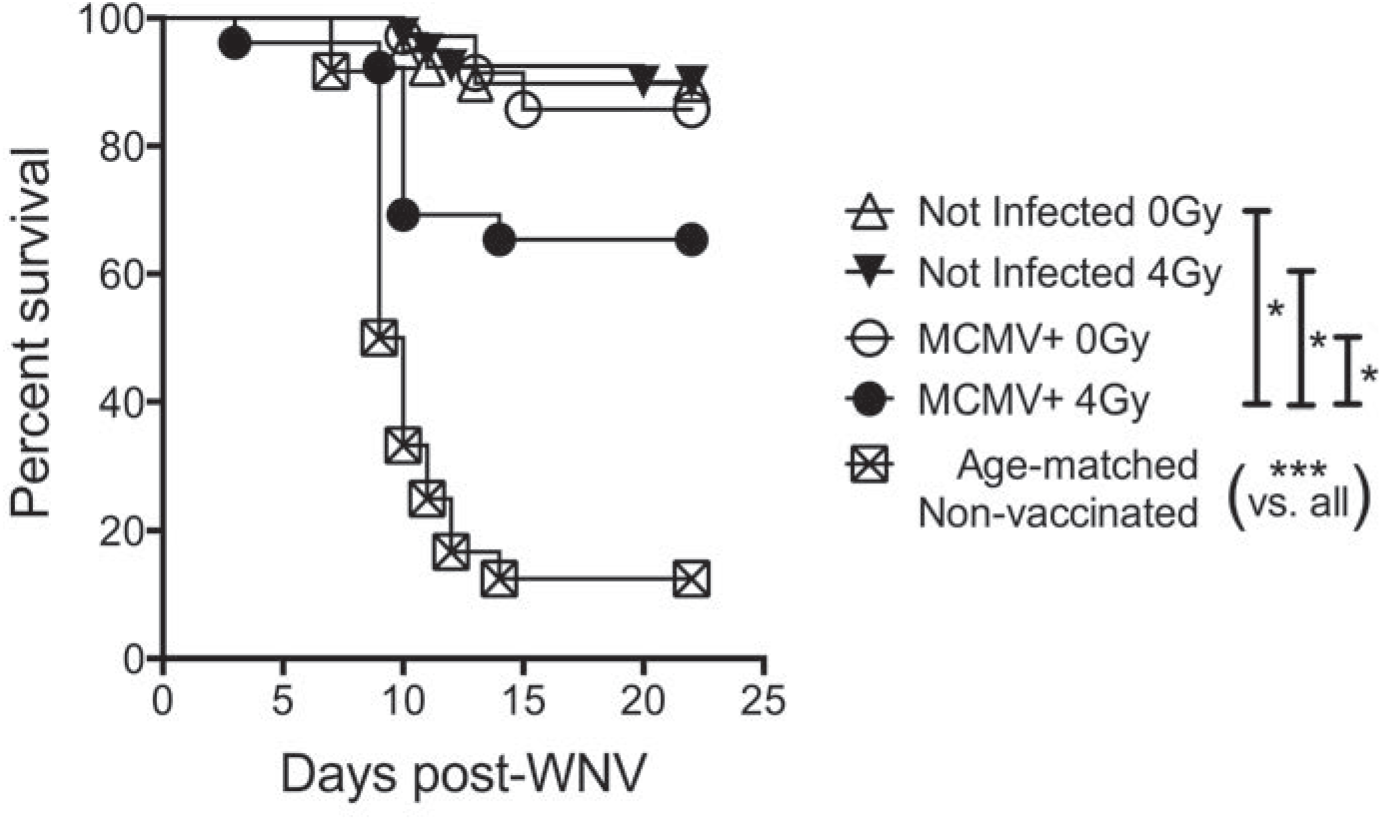
The additive effect of MCMV infection and WBI in youth on recall WNV survival in aged mice. Survival shown following RWN vaccination at approximately 19 months of age, and WNV challenge at approximately 21 months of age. Mice were infected with 2000 pfu WNV IP. Graph is a combination of both cohorts (no group statistically different between cohorts). Shown are results of individual comparisons from Kaplan-Meier tests. n ≥ 12 per group. Log-rank test for trend = ****.

### High-dose sub-lethal WBI in youth does not result in a loss of vaccination efficacy in old age

To investigate how WBI and CMV could damage the immune system, we followed immune cell populations throughout life (gated as in **Fig. S2**). Most populations were not significantly different across WBI doses by 19 months of age, at the time of vaccination, including B, NK-T and γδ (**Fig. S3**). While NK cell counts at 19 months appeared the most altered by WBI in youth, NK cells are dispensable for survival from WNV (37). NS4b is the dominant CD8 T cell epitope responding to WNV and the RWN vaccine (38). At 7 days post-vaccination, NS4B tetramer positive (NS4b+) CD8 T cells were equivalently abundant in groups receiving different WBI doses and also between MCMV(-) and MCMV(+) groups (**Fig. 3A, (39)**). Anti-Ki-67 antibody marks cells currently in the midst of any phase of cell cycle (G1,S,G2,M), but not those in interphase (G0) (40). Ki-67+ NS4b+ cells were also found in similar frequencies across all groups at the peak of the vaccination response (**Fig. 3B, (39)**). The exception were the MCMV(+) 4Gy mice, which exhibited lower representation of these cells compared to their 0Gy counterparts when data from cohorts were combined (**Fig. 3E**). NS4B+ cells also contained a similar proportion of Granzyme B^hi^ cells across all groups (**Fig. 3C, (39)**). In the memory phase (45 days post-vaccination and just prior to challenge), numbers of NS4B+ memory CD8+ T cells were comparable to adult vaccinated controls across all WBI and CMV groups (**Fig. 3F**). Finally, the analysis of sera at 45 days post-vaccination revealed similar levels of neutralizing of WNV-specific antibodies (as judged by the plaque reduction neutralizing titer (PRNT) assay), revealed that neither WBI dose in youth, nor life-long MCMV status significantly altered the neutralizing potential of serum antibody generated by RWN vaccination in old mice (**Fig. 3G, (39)**), albeit the titers were significantly lower compared to adult vaccinated animals **(Fig. 3G, (39))** confirming our published work (36).

**Figure 3:**
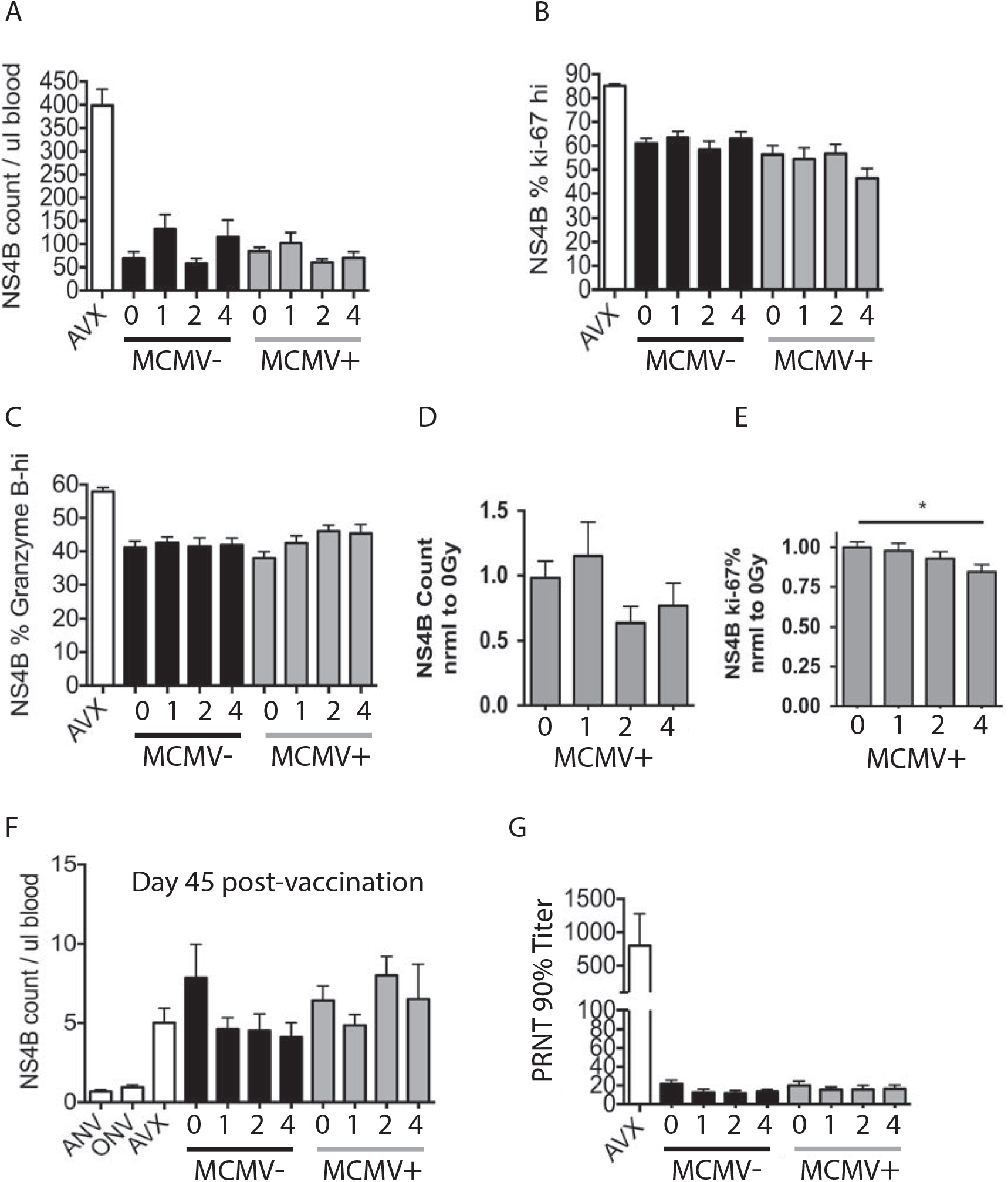
Vaccine response is equivalent in old mice exposed to WBI in youth. All mice injected with 10^5^ pfu RWN IP on the same day. AVX = adult vaccinated controls (5 months). All other groups ≥ 19 months. Numbers on x-axis indicate dose of WBI in youth in Gy. M indicates mice that have had MCMV for life. All data from PBMC. (A-D, H, I) shown from a single cohort, n ≥ 8 per group. (A) Counts of NS4B tet+ CD8 T cells on day 7 post-RWN. (B) Percent of Ki-67 hi, NS4B tet+ CD8 T cells on day 7 post-RWN. (C) Percent of NS4B tet+ CD8 T cells that are Granzyme B-hi on day 7 post-RWN. (D-E) data shown from combined cohorts normalized to 0Gy (No WBI) MCMV(+) mice in each group, n ≥ 15 per group. (D) Counts of NS4B+ CD8 T cells in MCMV+ mice on day 7 post-RWN. 1-way ANOVA = ns. (E) Percent of Ki-67-hi in MCMV(+) mice on day 7 post-RWN. 1-way ANOVA = ns. Results of Dunnet’s Multiple comparison shown. (F) Counts of established memory NS4B tet+ CD8 T cells on day 45 post-RWN. ANV = adult non-vaccinated controls, ONV = age-matched old nonvaccinated controls. All other labels as in (A). (G) Maximum dilution of serum collected 45 days post-RWN vaccination, that achieves 90% reduction of WNV plaques at 100 pfu.

To examine functional T cell responses, at 45 days post-vaccination, PBMCs were harvested and stimulated with WNV peptides recognized by CD8 T cells in the presence of Brefeldin A. Only slight, non-significant differences were noted across WBI doses and MCMV status with regard to individual cytokine production (**Fig. 4A, (39)**). In fact, polyfunctional cytokine production in response to WNV-specific peptides appeared more robust in the MCMV(+) 4Gy group (**Fig. 4B, (39)**). Finally, at the height of WNV challenge, 8-days post-WNV infection, NS4b+ populations responded with roughly equivalent size (**Fig. 4C–4D, (39)**), Granzyme B production (**Fig. 4E**), and proliferation (**Fig. 4F**) regardless of life-long MCMV, WBI or both in youth.

**Figure 4:**
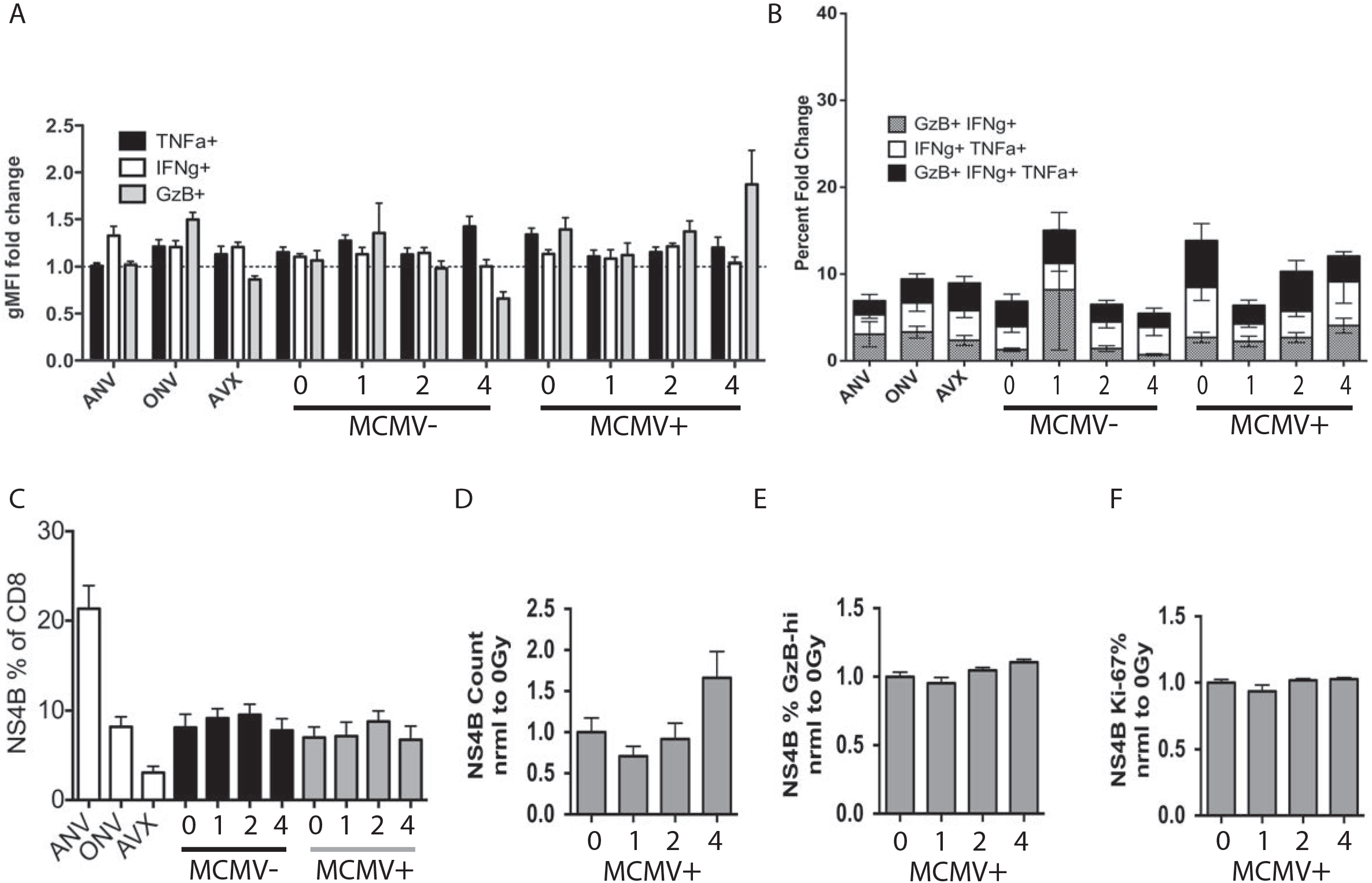
T cell function and responses to WNV challenge in mice exposed to WBI in youth. AVX = adult vaccinated controls, ANV= adult non-vaccinated controls (5 months). ONV= non-vaccinated mice, age-matched to cohort (21 months). Numbers on x-axis indicate dose of WBI in youth in Gy. M indicates mice that have had MCMV for life. (A-B) shown from a single cohort, n ≥ 10 per group, except ANV = 4. PBMC from mice on day 45 post-RWN vaccination were isolated and stimulated with CD8-specific WNV peptides for 5 hours in the presence of Brefeldin A. (A) Fold change (peptides / no peptides) of individual cytokine production by geometric MFI. (B) Fold change in the percentage of polyfunctional CD8 T cells (peptides / no peptides) of CD8 T cells treated as in (A). (C-D) data shown from a single cohort. (C) Percent of NS4B tet+ CD8 T cells in PBMC on day 8 post-WNV infection. (D-F) data shown from combined cohorts normalized to 0Gy (No WBI) MCMV(+) mice, all ANOVA post-tests = ns. (D) NS4B tet+ CD8 T cells in PBMC on day 8 post-WNV. (E) Percent of NS4B tet+ T cells that are Granzyme B-hi on day 8 post-WNV infection. (F) Percent of NS4B tet+ T cells that are Ki-67-hi on day 8 post-WNV infection.

### WNV infection causes MCMV reactivation

As expected, latent MCMV was reactivated by WBI in a dose-dependent manner, across a variety of tissues (41) (**Fig. S4A-D**), with the highest reactivation seen at 4Gy. Prolonged immunological marks of reactivation were seen as a significant increase in representation of m139+ CD8 T cells on d 30 post WBI between unirradiated, 2Gy and 4Gy group in an irradiation dose-dependent manner. Because primary and memory immunity to vaccination were not altered by WBI, but survival from WNV was worsened specifically in the MCMV(+) 4Gy group, we wanted to examine the potential interaction of WNV and MCMV co-infection. MCMV is kept latent by a combination of immune factors, including antibody, NK, and CD8 T cells (42). Low-level MCMV reactivation may be traceable through populations of MCMV-specific expanding CD8+ T cells (26). m139 and m38 tetramer positive (m139+ and m38+) CD8 T cells are two such MCMV-specific CD8+ populations that undergo life-long expansion in latently infected mice (43). We found that both m139+ and m38+ CD8+ T cells increased GzB production during the height of WNV infection (**Fig. 5A**). In order to rule out bystander activation or a WNV epitope mimetic, we serially infected young mice to create antigen-specific, pMHC tetramer-traceable bystander memory population (**Fig. 5B**). Mice were first infected with MCMV. After MCMV latency, mice were infected with Listeria monocytogenes genetically modified to express the SIINFEKL epitope (LM-Ova) and after that response matured into memory, mice were infected with WNV, and cells analyzed for effector responses by pMHC-Tet+GzB+ staining. As expected, GzB content was highest in NS4b+ (WNV-specific) T cells at the height of T cell response to WNV infection. However, GzB was significantly increased in m38+ and m139+ (MCMV-specific) T cells, but not in bystander Ova memory T cells, nor in the entire remainder of CD44-hi (memory pool) T cells (**Fig. 5C**). Finally, to directly establish MCMV reactivation during WNV co-infection, we collected liver in mice with latent MCMV on day 3 following WNV infection. MCMV+ genomic content by qPCR was approximately 10-fold higher in hepatocytes during WNV infection, compared to hepatocytes in mice that were mock-infected with WNV (**Fig. 5D**). Similar results were obtained with other stressors administered to CMV-positive mice, including corticosterone, L. monocytogenes infection and different doses of radiation, using sensitive primers to detect both MCMV DNA (replication) and gene expression (transcription) (**Fig. 5E**). Therefore, latent MCMV was readily reactivated by various stressors, including the WNV infection.

**Figure 5:**
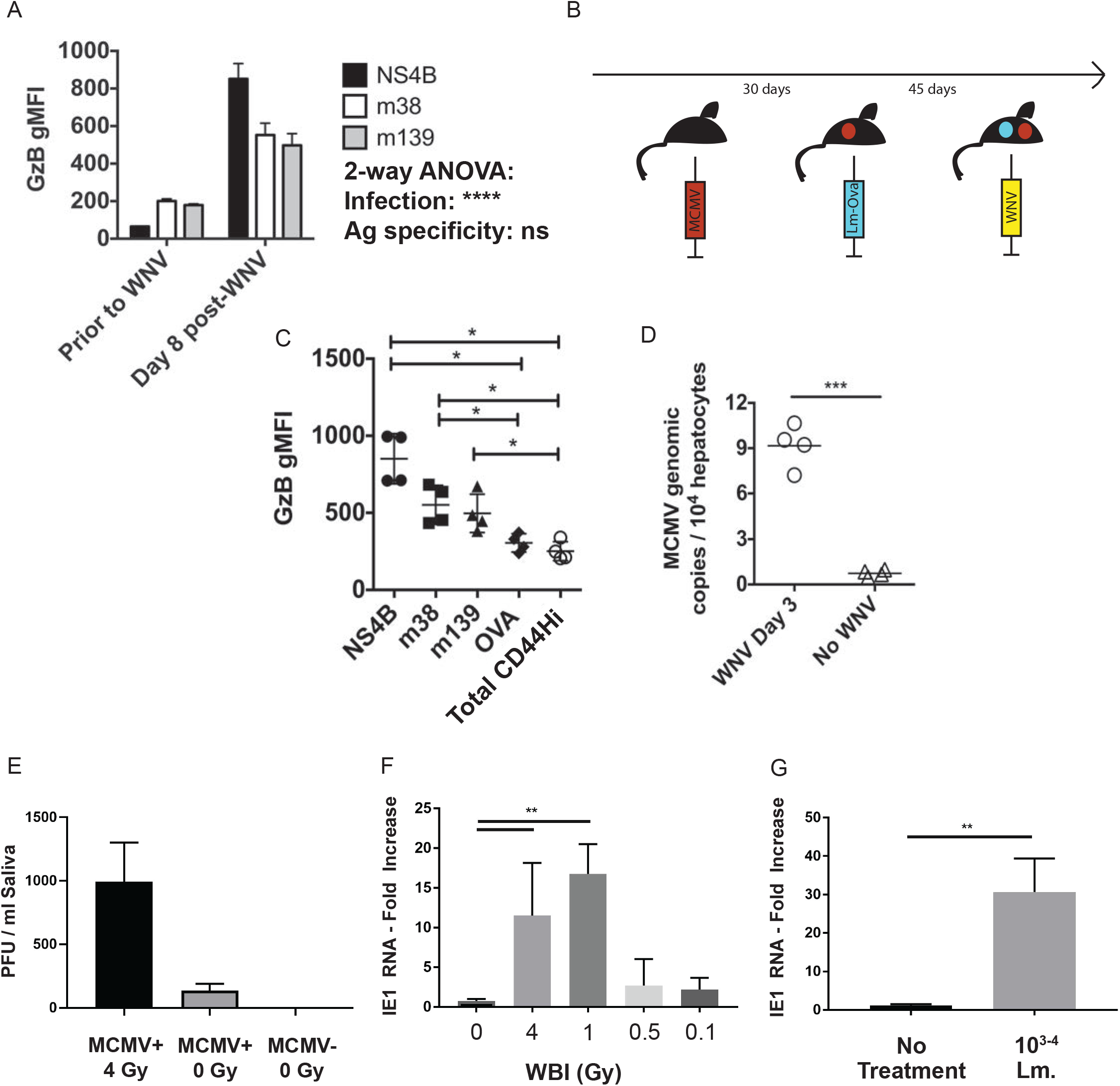
WNV infection causes reactivation of MCMV in adult mice. (A) Geometric MFI of Granzyme B in tetramer-specific CD8 T cell populations in PBMC from n=4 adult MCMV(+) (latent) mice prior to WNV infection, and on day 8 post-WNV infection. Tukey post-tests between tetramers = ns. (B) Experimental design for data shown in (C). Mice were infected IP with 200 pfu smith strain MCMV, then rested for 30 days. Mice were then infected IV with 10^5^ cfu Listeria monocytogenes expressing OVA epitope (LM-Ova). Memory populations were allowed to mature for 45 days. Mice were then infected with 2000 pfu WNV IP. (C) Geometric MFI of Granzyme B in tetramer-specific CD8 T cell populations from splenocytes collected on day 8 post-WNV infection from mice in experiment (B). CD44hi = the remainder of tetramer non-specific, CD44-hi T cells. Shown are the results of Bonferroni post-tests. (D) Genomic copies of MCMV from livers of mice with latent MCMV, either without WNV, or on day 3 post-WNV infection, from qPCR for MCMV genomic DNA. (E) MCMV genomes recovered from saliva day 3 post 4 gy irradiation and transcription of MCMV IE1 in the salivary gland 3 days after a titrating dose of irradiation (as indicated) or 10^3-4^ CFU of *Listeria monocytogenes* (Mann-Whitney).

### MCMV-specific immunity is eroded in old mice that harbored latent MCMV during high-dose, sub-lethal WBI in youth

Given the propensity of WBI to cause MCMV-reactivation (41) (**Fig. S4 & 5D,E**), we wanted to examine MCMV-specific immunity following WBI and repopulation. At 13 months of age, groups of mice were sacrificed and splenocytes were subjected to peptide stimulation with m139 peptide in the presence of Brefeldin A. CD8 T cells from mice who received 4Gy of WBI in youth produced lower amounts each of Granzyme B, IFNγ, and TNFα following peptide stimulation, compared to other, lower WBI doses (**Fig. 6A,B**). MCMV(+) 4Gy mice further began to display increased intensity of PD-1, an activation/exhaustion marker, on m139+ CD-8 T cells by 13 months of age (**Fig. 6C**). By 19 months of age, WBI in youth resulted in a suppressed MCMV-specific serum Ab production (**Fig. 6D**). CD127 expression is known to decrease in exhausted populations of CD8 T cells in some chronic infections (44, 45). We found that WBI in youth decreased CD127 expression in a dose-dependent manner in m139+ CD8 T cells by 19 months of age (**Fig. 6E**). KLRG1 is a marker whose expression on memory CD4 T cells correlates with replicative senescence (46). The KLRG1-high portion of memory CD4 T cells increased significantly in MCMV(+) 4Gy group by 19 months of age (**Fig. 6F**). Neither marker of senescence/exhaustion was apparent in MCMV(-) mice regardless of WBI dose in youth (**Fig. 6** and **S5A–S4B**), strongly implying that these effects were mediated by MCMV reactivation, not WBI alone. To examine whether lasting DNA damage could be behind these effects, we analyzed levels of γH2AX in CD8+ T cells, a marker that labels double-strand DNA breaks currently under repair. We found no increase in standing DNA damage in any group at 13 months (**Fig. 6G, S5C**), implying that the signs of exhausted/senescent anti-CMV response were not mediated by lasting WBI-induced DNA damage. Moreover, at a steady state, 8 months post irradiation (13 months of life) there were no signs of increased MCMV DNA replication, suggesting stable latency regardless of prior irradiation in youth (**Fig. S5D**).

**Figure 6:**
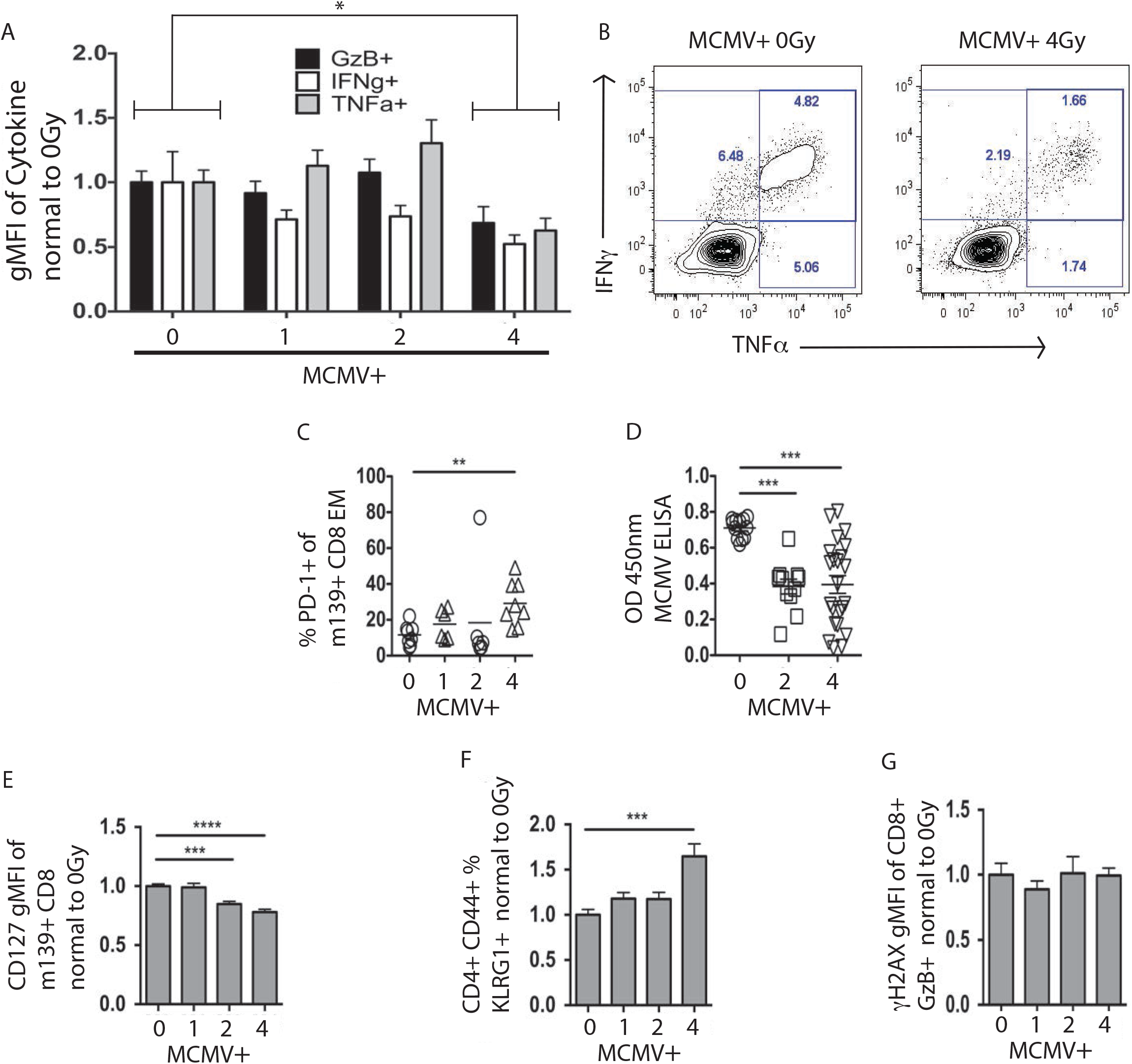
High-dose sub-lethal WBI in youth in MCMV+ mice taxes MCMV-specific immunity. (A-B) Splenocytes from MCMV+ mice at 13 months of age were stimulated for 6 hours with m139 peptide in the presence of BFA. (A) Percent of individual cytokines normalized to 0Gy (No WBI) MCMV+ group. 2-way ANOVA post-tests M0 vs M4 = ***. (B) Representative flow plots, as in (A). (C) Percent of PD-1+ cells from PBMC at 13 months of age. Statistical outlier shown but not included in statistical analysis. (D) Results of MCMV ELISA from plasma collected at 19 months of age. (E) m139+ CD127 MFI of PBMC at 19 months of age from combined cohorts, normalized to 0Gy (No WBI) MCMV+ mice. (F) KLRG1-hi percent of memory CD4 T cells from PBMC at 19 months of age from combined cohorts, normalized to 0Gy (No WBI) MCMV+ mice. (G) Gamma-H2AX MFI of GzB+ CD8+ splenocytes at 13 months of age, normalized to 0Gy (No WBI) MCMV+ mice. (C-G) Results of Dunnet’s post-tests shown.

### WNV-mediated CMV reactivation critically predicts lethal outcome

We examined proliferation and differentiation of m139-specific CD8+ T cell populations during RWN vaccination and WNV challenge in mice that have received different doses of WBI in youth. Regardless of radiation dose, m139-specific populations exhibited similar (8-17%) fraction of dividing (Ki-67+) cells at 19 months of life, 7-days post-RWN vaccination, and 60 days post-RWN vaccination (**Fig. 7A**). However, m139+ populations were significantly more prolific (28-35% Ki-67+) in old mice on day-8 post WNV challenge regardless of prior irradiation (**Fig. 7A**), indicating that MCMV reactivation also occurred in old vaccinated mice during WNV infection, and that it was relatively independent of irradiation dose in youth. That was consistent with increased GzB production, which was enhanced twofold in the m139+ population during WNV challenge across WBI doses, and disproportionately so (threefold, p<0.001) in the group MCMV(+) 4Gy, which exhibited decreased survival (**Fig. 7B**). Upon detailed examination of the MCMV(+) 4Gy group, we could precisely stratify it by GzB levels in m139-specific CD8 T cells into high and low expressors; those mice that perished from WNV challenge had significantly higher levels (> 1500 relative MFI; **Fig. 7C**) of m139+ cells during the WNV response, compared with those that survived. By contrast, no parameters of anti-WNV immunity correlated with death in the MCMV+ 4Gy group (see **Fig. 6**). Rather, mice that perished from WNV appeared to have enhanced WNV-specific T cell responses, though WNV-specific antibody was equivalently abundant and neutralizing (**Fig. 5E-F**).

**Figure 7:**
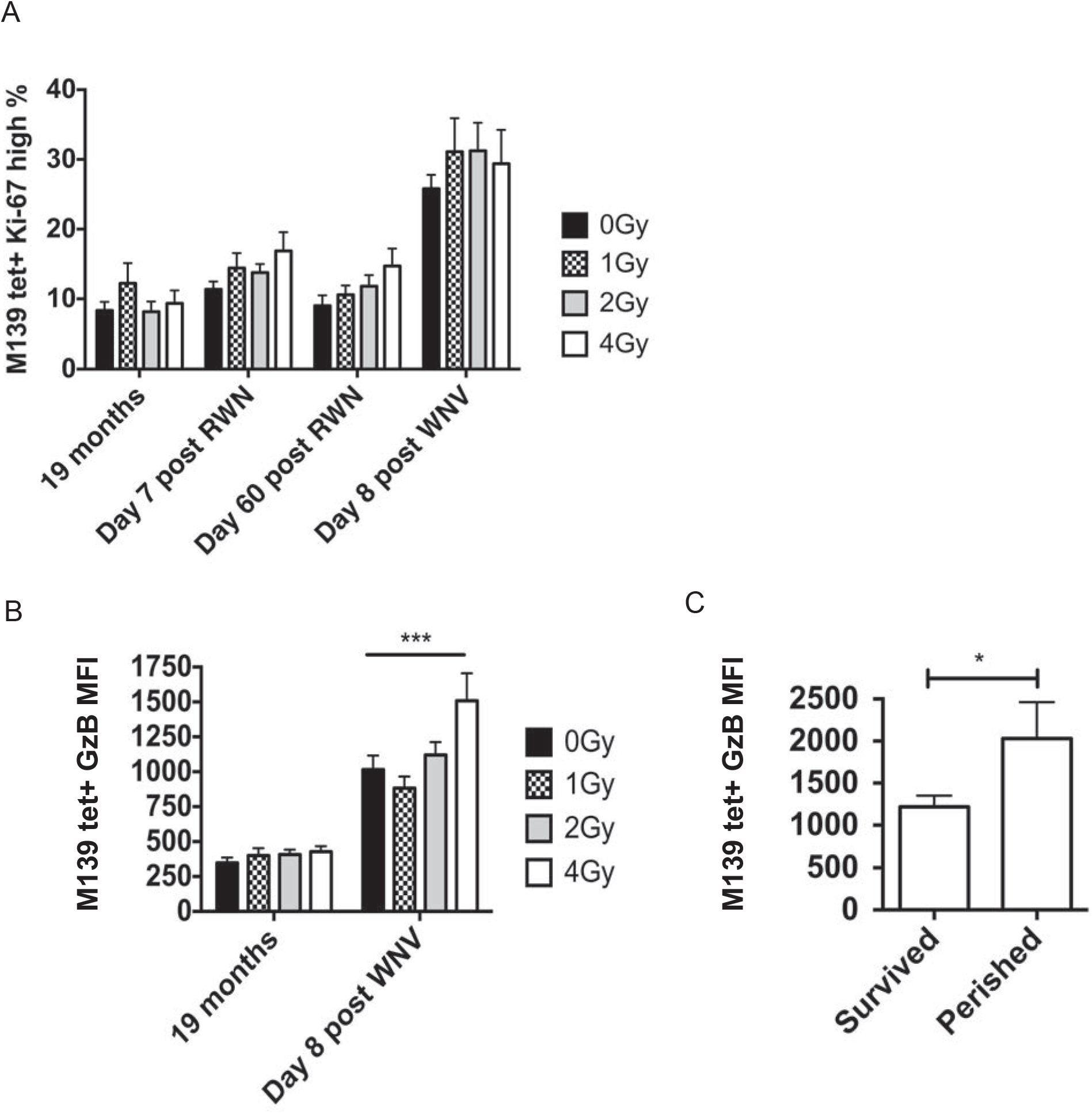
MCMV reactivation occurs in old mice during WNV infection, and is associated with increased mortality. Numbers on x-axis indicate dose of WBI in youth in Gy. M indicates mice that have had MCMV for life. (A) Percentage of Ki-67 hi CD8 T cells within the m139 tetramer-specific population, at various time points in late life. (B) geometric Granzyme B MFI within the m139 tetramer-specific population at 19 months or on day 8 post-WNV infection. ANOVA of 19 months = ns. Shown is the result of Bonferroni post-test between radiation groups. (C) Geometric MFI of Granzyme B in mice from the MCMV(+) 4Gy group, stratified between mice that survived and mice that perished from WNV challenge.

Overall, our data supports a model in which MCMV reactivation during WNV co-infection increases the likelihood of lethal outcome. Two lines of evidence support this conclusion: (i) MCMV-specific immunity was functionally defective in the MCMV(+) 4Gy group (**Fig. 6**), allowing for enhanced MCMV reactivation and inflammation during WNV infection, ultimately leading to increased mortality from co-infection by mechanisms that are under investigation; (ii) by contrast, WNV-specific immunity was functionally equivalent regardless of CMV status or radiation dose (**Figs. 3–5**). These results are discussed below.

## DISCUSSION

In our prior work (Pugh et al., Aging Cell), we found no evidence that a single exposure to ionizing radiation of up to 4Gy in youth can cause immune defects in the old age (REF. Pugh, Aging Cell). To further address this issue, we have modeled a frequent human persistent herpesvirus infection with cytomegalovirus in mice, and examined whether ionizing radiation event in youth would modulate immunity in the presence of MCMV. We found that following WNV vaccination and challenge, old mice with MCMV that received 4Gy irradiation in youth exhibited worse survival than any other group of animals, whereas neither high-dose WBI or MCMV alone made either the survival or the immune responses any worse compared to aging alone.

To elucidate the basis behind this surprising relationship, we carefully examined WNV immunity and found no differences between groups with high and low mortality in humoral or cellular immune responses. Because increased mortality was exclusive to the MCMV(+) 4Gy group, we reasoned that an MCMV-specific immune defect, linked to high WBI, would be a likely culprit. It was possible that 4Gy WBI generally weakened memory cells. However, memory T cells in MCMV(-) 4Gy mice did not display any markers of exhaustion or senescence following repopulation. Therefore, it was unlikely that repopulation stress alone caused MCMV-specific immune defects. We next suspected that WBI may have triggered MCMV reactivation, and found that MCMV reactivated in a WBI-dose dependent manner, such that the reactivation of MCMV was of greater magnitude and involves a broader array of tissues in 4Gy-exposed mice compared to 2Gy-exposed mice. Steady-state levels of latent MCMV were no higher in 4Gy mice than in 0Gy mice following repopulation, and therefore the MCMV burden was not permanently increased by WBI and reactivation. We further found that irradiated MCMV+ mice exhibited dose-dependent alterations in MCMV-specific T cell responses, which were most pronounced in 4Gy-irradiated mice. Together with the finding that WNV infection also caused MCMV reactivation, these results suggested that the most likely scenario is that WBI weakened MCMV immunity through the systemic MCMV reactivation event following WBI, and that lifelong subclinical reactivations further potentiated this effect. Upon WNV infection, another MCMV reactivation challenged the MCMV response already weakened by prior irradiation, making animals susceptible to succumbing to a combination of WNV and MCMV. Further work is necessary to elucidate whether systemic reactivation without WBI is sufficient to weaken MCMV immunity, and whether memory T cells are more susceptible to activation-induced exhaustion following WBI and repopulation.

The danger of co-infection with MCMV and WNV is at first counter-intuitive, in that Th1-mediated responses to either virus should aid in clearing the other. Following this logic, latent gamma-herpesvirus has been shown to be protective in the context of bacterial co-infection in adult mice (47), and we have also shown that latent MCMV is protective in Listeria infection in old mice by broadening TCR repertoires (Smithey, PNAS). The increased mortality due to MCMV co-infection is therefore specific to old mice. MCMV utilizes a variety of immune evasion mechanisms, however, we did not see evidence of reduced immune responsiveness in mice that died in our challenge experiments – in fact, we observed increased signs of immune reactivity against CMV and WNV, which is not consistent with immune evasion. If mortality from WNV is immunopathogenic in old mice, then increased immune activation from MCMV might increase mortality. However, WNV immune infiltrates in the brains of old immunocompetent mice are few compared to other immunopathogenic infections (48) making this scenario unlikely. It is possible that CMV could increasingly infect the CNS as a result of increased inflammation and WNV entry into the brain during co-infection. The precise mechanism of MCMV and WNV co-infectious mortality remains to be elucidate and experiments are in progress to address it conclusively.

Latent CMV can be found in a variety of tissues in both infected mice and humans (41, 49). CMV reactivation can occur in association with cell differentiation, immune compromise, or tissue damage, and has recently been reported during coinfection in a model of herpesvirus and helminth infection (50). The pathology of WNV includes viremia and infection of a variety of organs (51). It is therefore not surprising that WNV could reactivate CMV during the course of infection. We have confirmed that similar MCMV reactivation occurs in mouse models of Listeria infection and steroid treatment. It remains to be seen what infectious burden if any is sufficient to reactivate CMV in humans. Our work points to the possibility of additional risks borne by older CMV+ individuals during systemic infections due to acute CMV reactivation. This also implies that the enhanced mortality associated with CMV is perhaps due to the cumulative effect of opportunistic CMV reactivation events during new infections throughout life, rather than the static immune burden of latent CMV. Individual history of systemic infection may therefore be an important predictor of mortality in CMV+ individuals.

Our results also imply that the trajectory of immune health in individuals exposed to WBI is highly dependent on CMV status at the time of exposure. Because CMV is highly prevalent in human populations, radiobiology studies should discriminate based on latent pathogen status, and radiobiology studies with animal models should include latent infections that closely mirror human populations.

## METHODS

### Mice

Adult (< 6 month) Male C57BL/6 mice were acquired from Jackson Laboratories and housed under specific pathogen-free conditions in the animal facility at the University of Arizona (UA). All experiments were conducted by guidelines set by the UA Institutional Animal Care and Use Committee. As needed, mice were euthanized by isofluorane and spleen was collected into complete RPMI supplemented with 5 or 10% Fetal Bovine Serum (FBS). Blood was taken from the heart or retro-orbitally for cross-sectional and longitudinal harvests, respectively and red blood cells were hypotonically lysed.

### Viruses and Vaccine

Smith Strain MCMV was obtained as multiple passage stock from Drs Ann Hill (Oregon Health & Science University, Portland, OR) or as a low-passage infectious clone from and Wayne Yokoyama (Washington University, St. Louis, MO). Animals were injected IP at 10^5^ pfu / mouse, and both stocks showed identical acute responses. West Nile Virus (WNV): Strain 385-99, a kind gift from Robert Tesh, was injected IP at 2000 pfu / mouse. Replivax WNV (kind gift from Drs P. Mason, N. Bourne and G. Milligan, U. Texas Med. Branch, Galveston, TX) was injected IP at 10^5^ pfu / mouse. Verification of viral titer and production of RWN stock are described elsewhere (36).

### Peptide Stimulation

Blood samples were taken at 45 days following RWN vaccination, hypotonically lysed, and stimulated ex-vivo with a pool of NS4b 2488-2496 and E 347-354 peptides (21st century Biochemicals, Marlborough MA) both at 10^-6^ M. Stimulation took place over 6 hours in the presence of BFA. Splenocytes from 13 month-old mice during cross-sectional harvest were treated identically for 5 hours with 2 ng / μl MCMV m139 peptide.

### MCMV ELISA

We used commercial MCMV ELISA kits (Cat# IM-811C, XpressBio, Thermont, MD). Serum samples were applied at 1:50, 1:250, and 1:500 dilutions in duplicate, and at 1:50 in wells coated with uninfected cell lysate (negative control).

MCMV-mouse serum was used as negative biological control. Serum AB was detected with anti-mouse HRP-bound AB in an enzymatic reaction, and OD was read at 450nm. Control wells for each mouse were subtracted from OD.

### Plaque Reduction Neutralization Test (PRNT)

Serial dilutions of mouse serum (1:10 minimum) were incubated with 100 pfu / well live WNV from the same stock received by mice, in a 96 well format, for 6 hours at 4C. Samples were then applied to a monolayer of Vero cells also in 96 well format, and allowed to incubate at 37°C with 5% CO_2_ for 25 hours. Resulting monolayers were fixed with ice cold 50% acetone 50% methanol for 30 minutes at −20C, and allowed to dry overnight. Resulting monolayers were assayed with anti-WNV antibody clone EG16, a kind gift from Michael Diamond, followed by peroxidase labeled Goat Anti-mouse IgG (XPL, inc, Gaithersburg MD). Infectious lesions were visualized in a DAB reaction. The dilution factor necessary for 90% reduction of infectious lesions was established by hand count. The average of duplicate assays per mouse was used.

### Irradiation

Whole-Body Irradiation was performed on a Gammacell Cs^137^ source irradiator calibrated by in-house physicist from the UA Health Sciences Center. Dosage was verified with thermal luminescence dosimeters (TLD) (Landauer Inc., Glenwood, IL) and TLDs from the Medical Radiation Research Center at the University of Wisconsin. Dosages fell within 5% of expected values. Effective dose rate ranged 70.2-68.36 cGy/min depending on the age of the source, and distance. For whole body irradiation a maximum of 8 mice were placed in sterile RadDisks (Braintree Scientific, Braintree, MA) with no separation. WBI occurred before noon on a light-dark cycle 7am-7pm.

### Flow Cytometry

Prior to each collection, voltages were manually calibrated to a common template using Rainbow Beads (BD Biosciences, San Jose, CA), to insure accurate MFI tracking over time. Fluorescent conjugated α-Mouse antibodies against CD3(SK7), CD4(MCD0430), CD8a(S3-6.7), CD62L(MEL-14), CD44(IM7), α-Ki-67(B56), CD127(A7R34), KLRG1(2FI), CD86(GL-1), B220(RM2630), NK1.1(PK136), CD49b(DX5), CD19(RM7717), IgM(II/41), MHC-ii(M5/114.15.2), were purchased from commercial sources. Tetramers against NS4b (H-2D(b) – SSVWNATTA), m139 (H2-K(b) – TVYGFCLL), m38 (H-2K(b) – SSPPMFRV) and Ova (H-2K(b) – SIINFEKL) were obtained from the National Institutes of Health Tetramer Core Facility. Staining occurred at 4C followed by fixation and permeabilization (FoxP3 kit, eBioscience). Blood and spleen counts occurred on a Hemavet cell counter (Drew Scientific, Dallas, TX). Samples were run on a Fortessa Flow Cytometer equipped with 4 lasers and using DiVa software (BD Biosciences). Compensation and analysis was performed using FlowJo software (Tree Star, Ashland, OR).

### qPCR

Tissues were harvested and placed into Eppendorf tubes with 1mL Tri-Reagent (Life Technologies/Ambion, Grand Island, NY) and sterile 1mm silica beads (Biospec Products, Bartlesville, OK), and immediately frozen in a dry-ice and 100% ethanol bath. Samples were thawed, bead-beaten, and DNA was extracted with phenol-chloroform. In the case of liver samples, DNA was subjected to two consecutive rounds of phenolchloroform extraction. Samples were normalized for DNA content and subjected to qPCR in quadruplicate with SYBR Green master mix (Life Technologies). Primers for either MCMV IE1 (*IE1-1*: CCC TCT CCT AAC TCT CCC TTT, *IE1-2*: TGG TGC TCT TTT CCC GTG) or C57BL/6 beta-actin (*BA-1*: AGC TCA TTG TAG AAG GTG TGG, *BA-2*: GGT GGG AAT GGG TCA GAA G) were used. Serial dilutions of plasmids (pCR-Blunt) with either IE1 or beta-actin insertion sequences were used in each plate to establish real counts and primer efficiencies. Primers and plasmids were developed by Bijal Parikh, and plasmids were supplied as a kind gift from Wayne Yokoyama. Samples were run on ABI 7900 (Life Technologies/Applied Biosystems) at the University of Arizona Genomics Core facility. Alternatively, tissues were harvested from mCMV infected and uninfected age matched controls and collected into microcentrifuge tubes containing 5x volumes RNAlater (Millipore Sigma) and stored at −80. Each sample was thawed and processed using Nucleospin RNA plus with DNA removal (Machery-Nagel) preps per manufacturer’s protocol. Reverse transcription was performed using OmniScript reverse transcriptase (Qiagen) and oligo-dT primers. qPCR to measure the expression *β-actin* and *IE1* of was performed using PowerUP SYBR Green Master Mix (Applied Biosciences) on a Step One real-time PCR system (Applied Biosciences) using the following cycle protocol: an initial step at 2 min 50°C followed by 95° for 10 min, followed by 40 cycles of 95° for 15 sec, 60° for 1 min. Sample RNA content was normalized to *β-actin* and expression of *IE1* was compared using the 2^-ΔΔCT^ method. Cycle 32 was set as a negative cut-off based on uninfected controls. Primer sets were gifted by Chris Benedict, PhD, La Jolla Institute of Immunology.

### Statistics

Statistical analysis was performed using Prism 6.0 (Graphpad Software). When data from multiple cohorts were combined for analysis, data from each cohort were normalized to the average of M0 group from that same cohort, to avoid cohort-specific biases. Significance is noted as follows throughout: ns = not significant, **** = p<0.0001, *** = p<0.001, ** = p<0.01, * = p<0.05. All error bars shown are SEM.

### Data

Data comprising all of the main figures is available in Supplementary Data 6.

## ACKNOWLEDGEMENTS

The authors wish to thank Drs. Michael Diamond, Wayne Yokoyama (Washington University), and Ann Hill (OHSU) for reagents and protocols. Thanks are also due to Drs. Giovanni Bosco (Dartmouth College), Ted Weinert, Kirsten Limesand, Jeff Frelinger and Mike Kuhns (University of Arizona) for critical input and suggestions. Special thanks to Dr. Wendell Lutz (Univ. of Arizona Cancer Center) for radiation dosage calibration and instruction, and Richard Wagner (University of Arizona) for help with radiation safety.

## CONFLICT OF INTEREST STATEMENT

The authors have no conflicts of interest to declare.

## SUPPORT

Supported in part by the USPHS awards AG020719 and AG048021 and the contract from the National Institute of Allergy and Infectious Diseases HHSN272200900059C, to the Radiation Effects Research Foundation (Dr Nakachi, PI) and its subcontract to Dr. Nikolich-Zugich. The Radiation Effects Research Foundation (RERF), Hiroshima and Nagasaki, Japan, is a private, non-profit foundation funded by the Japanese Ministry of Health, Labor and Welfare and the United States of America Department of Energy (US-DOE), the latter in part through DOE Award DE-HS0000031 to the US National Academy of Sciences. This study was based on RERF Research Protocol RP#4-09 and was supported by the US National Institutes of Health (NIAID Contract HHSN272200900059C). The views of the authors do not necessarily reflect those of the two governments.

## AUTHOR CONTRIBUTIONS

JLP designed and performed experiments, and wrote the manuscript; CPC performed experiments; ASS performed experiments; JLU performed experiments and designed assays; JP-T performed experiments; T.H. and K.N. provided critical advice; JN-Z designed experiments, directed the study, wrote and edited manuscript.

## DATA AVAILABILITY

Data comprising all main figures is included in Supplementary Data file 6.

## ABBREVIATIONS

WBI: Whole Body Irradiation
MCMV: Murine Cytomegalovirus
Gy: Gray (unit of radiation)
RWN: Replivax West Nile
WNV: West Nile Virus

---

**Figure S1.**
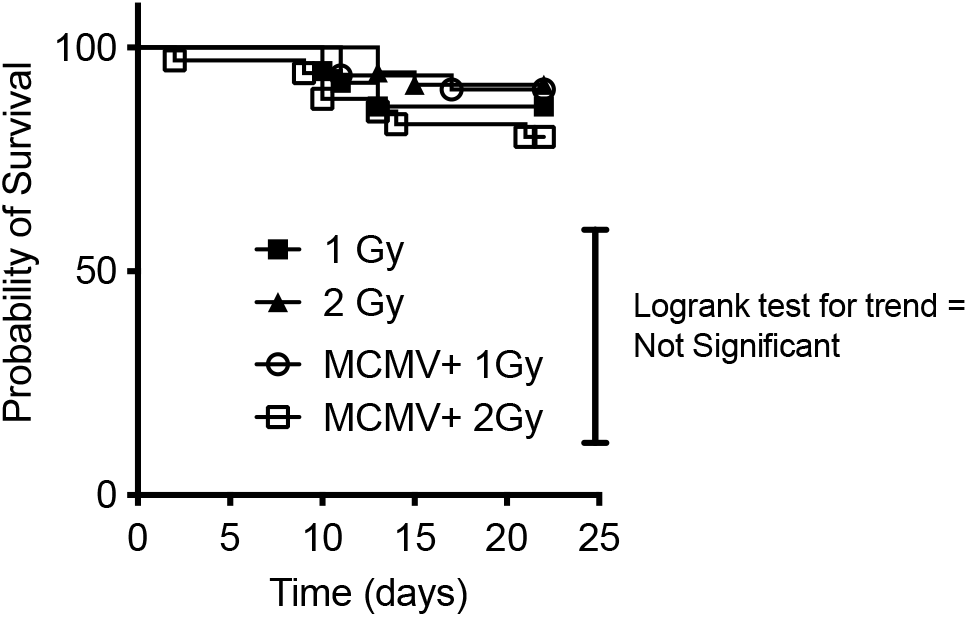
1 or 2 Gy WBI in youth, alone or in combi-nation with MCMV infection, does not significantly effect survival from WNV challenge in old age. Survival shown following RWN vaccination at approximately 19 months of age, and WNV challenge at approximately 21 months of age. Mice were infected with 2000 pfu WNV IP. Graph is a combination of both cohorts (no group statistically different between cohorts). Groups of mice infected or not with MCMV, and or irradiated with 1 or 2 Gy WBI are not significantly different, nor is there a significant

**Figure S2:**
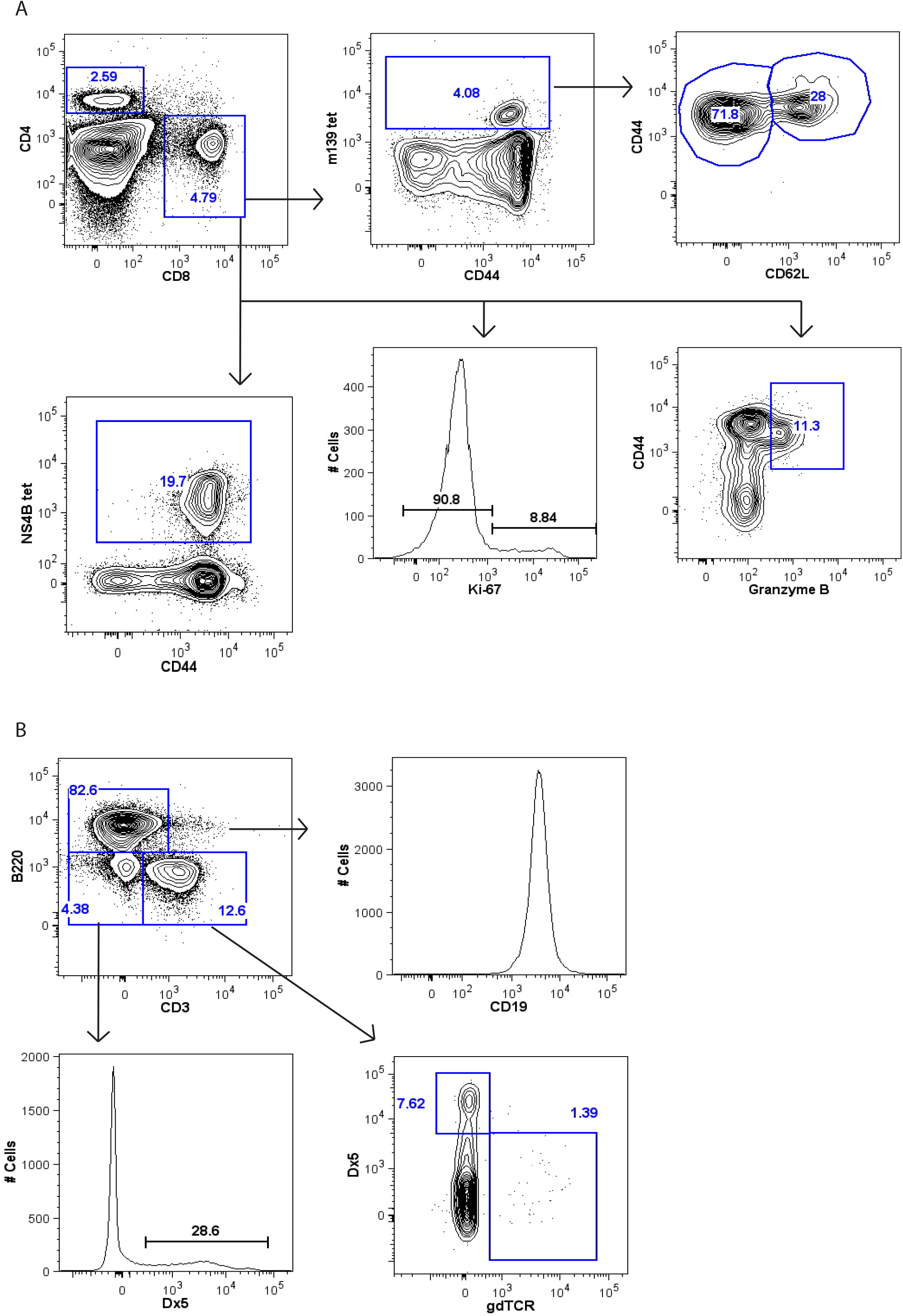
Gating strategies for flow cytometry. (A) Representative gating strategy for T cells and Tetramer-specific populations. (B) Representative gating strategy for B, NK, γδT and NKT cells.

**Figure S3:**
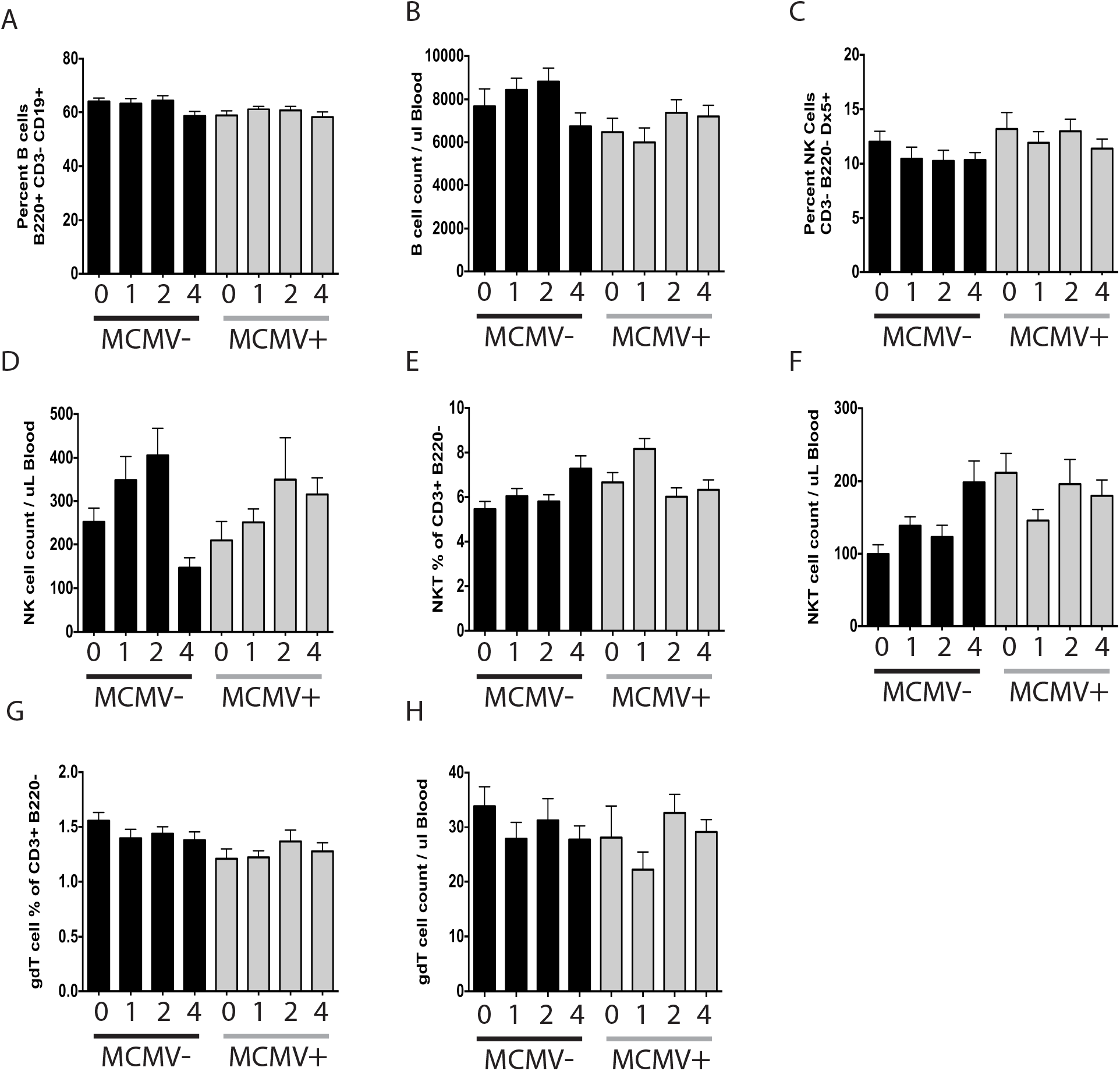
Standing counts and proportions of cells in PBMC of combined cohorts at 19 months of age. M indicates mice that have been infected with MCMV for life. Numbers indicate amout of WBI received in youth in Gy. (A) Proportion of B cells. (B) Count of B cells. (C) Proportion of NK cells. (D) Count of NK cells. (E) Proportion of NKT cells. (F) Count of NKT cells. (G) Proportion of γδT cells. (H) Count of γδT cells.

**Figure S4:**
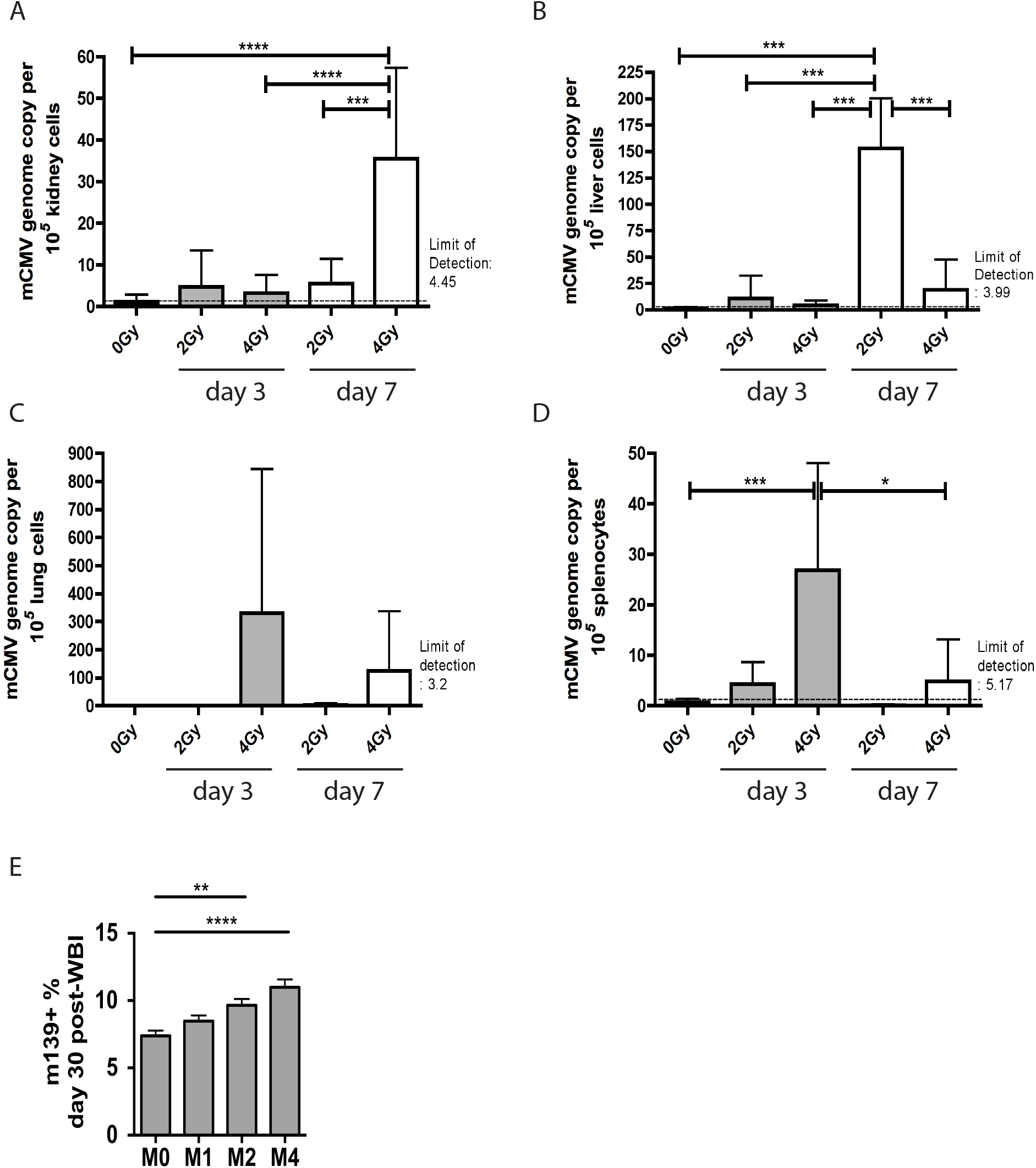
WBI reactivates MCMV. MCMV DNA assayed by qPCR from various tissues, normalized to signal from mouse Actin gene. All mice were adult C57BL/6 male mice subjected to WBI (2Gy or 4Gy) or no WBI (0Gy) 60+ days post-MCMV infection (latent MCMV). Tissues were harvested either 3 days or 7 days post-WBI. (A) Increase in MCMV genome copies in the kidney. (B) Increase in MCMV genome copies in the liver. (C) Increase in MCMV genome copies in the lung. (D) Increase in MCMV genome copies in the spleen. Shown are the results of Bonferroni post-tests. (E) m139+ proportion of CD8 T cells from PBMC of MCMV+ mice 30 days post-WBI. Shown are the results of Dunnet’s post-test.

**Figure S5:**
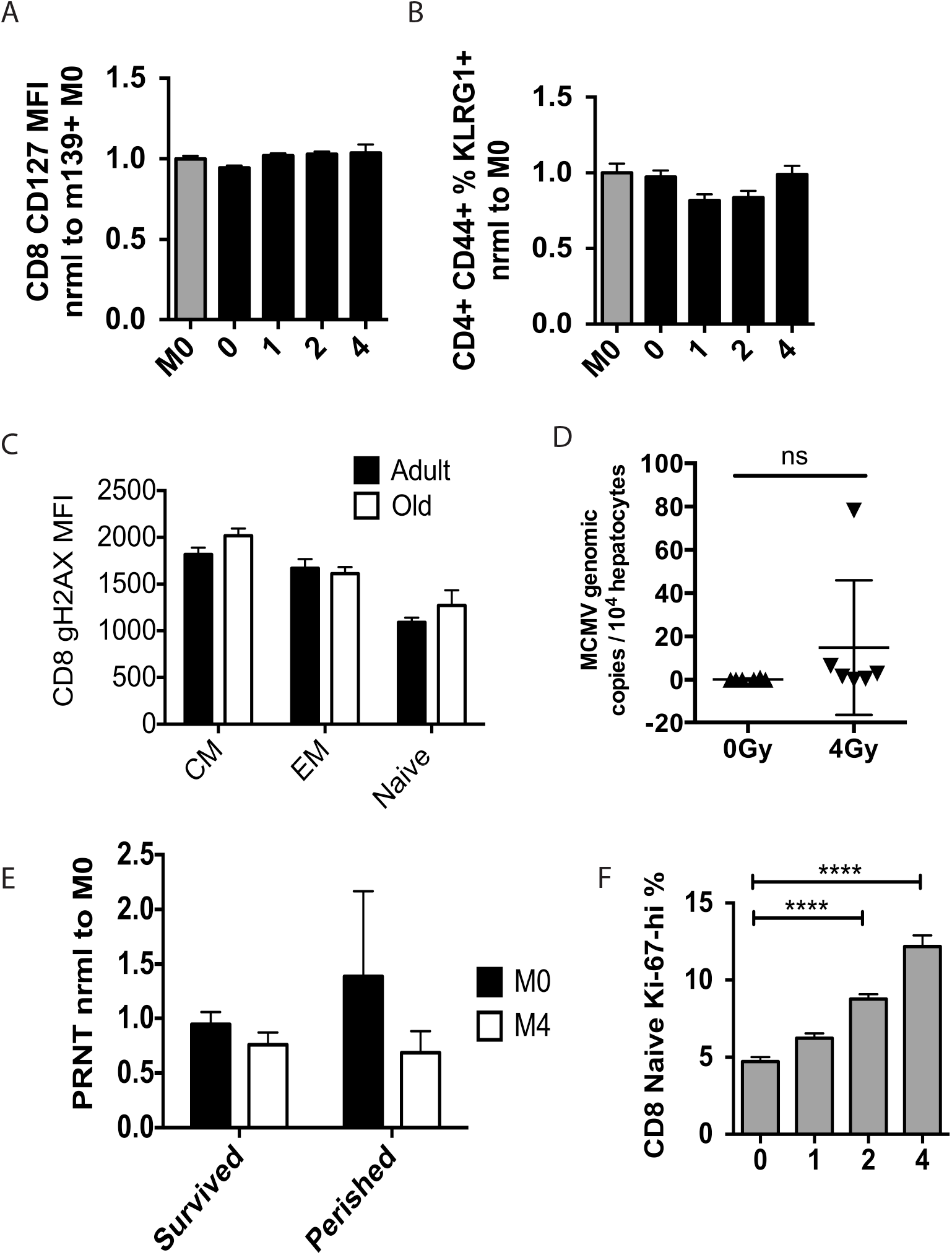
WBI alone does not increase senescence markers or latent MCMV burden, aging does not increase standing DNA damage in immune cells, and PRNT doesn’t correlate with death from WNV. M0 = MCMV(+) 0Gy, M4 = MCMV(+) 4Gy. (A) CD127 MFI in CD8 T cells of mice at 19 months of age from PBMC of combined cohorts, normalized to M0 group. (B) KLRG1+ proportion of memory (CD44-hi) CD4 T cells at 19 months of age from PBMC of combined cohorts, normalized to M0 group. (C) γH2AX MFI of adult (~5 months) and old (>18 months) male C57BL/6 mice in CD8 T cells from spleen. n=8 per age group. (D) MCMV genomic copies in hepatocytes at 13 months of age from mice with life-long MCMV subjected to WBI at 5 months age. Results of Mann-Whitney test shown (E) PRNT 90% titer reduction normalized to M0 survivors. Statistical post-tests are non-significant. (F) CD8 Naive Ki-67 hi portions in MCMV(+) mice from PBMCs 30 days post-WBI.

## Notes

### Competing Interest Statement

The authors have declared no competing interest.

